# The CarSR Two-Component System Directly Controls *radD* Expression as a Global Regulator that Senses Bacterial Coaggregation in *Fusobacterium nucleatum*

**DOI:** 10.1101/2024.12.13.628403

**Authors:** G C Bibek, Chenggang Wu

## Abstract

Two-component systems (TCS) enable bacteria to sense and respond to environmental signals, facilitating rapid adaptation. *Fusobacterium nucleatum*, a key oral pathobiont, employs the CarSR TCS to modulate coaggregation with various Gram-positive partners by regulating the expression of *radD*, encoding a surface adhesion protein, as revealed by RNA-Seq analysis. However, the direct regulation of the *radD*-containing operon (*radABCD*) by the response regulator CarR, the broader CarR regulon, and the signals sensed by this system remain unclear. In this study, chromatin immunoprecipitation followed by high-throughput DNA sequencing (ChIP-seq) identified approximately 161 CarR-enriched loci across the genome and a 17-bp consensus motif that likely serves as the CarR binding site. Notably, one such binding motif was found in the promoter region of the *radABCD* operon. The interaction of CarR with this binding motif was further validated using electrophoretic mobility shift assays (EMSA), mutagenesis, and DNase I footprinting analyses. Beyond regulating *radABCD*, CarR directly controls genes involved in fructose and amino acid (cysteine, glutamate, lysine) utilization, underscoring its role as a global regulator in *F. nucleatum*. Lastly, we discovered that RadD-mediated coaggregation enhances *radD* expression, and deletion of *carS* abolished this enhancement, suggesting that coaggregation itself serves as a signal sensed by this TCS. These findings provide new insights into the CarR regulon and the regulation of RadD, elucidating the ecological and pathogenic roles of *F. nucleatum* in dental plaque formation and disease processes.

**IMPORTANCE:** *Fusobacterium nucleatum* is an essential member of oral biofilms, acting as a bridging organism that connects early and late colonizers, thus driving dental plaque formation. Its remarkable ability to aggregate with diverse bacterial partners is central to its ecological success, yet the mechanisms it senses and responds to these interactions remain poorly understood. This study identifies the CarSR two-component system as a direct regulator of RadD, the primary adhesin mediating coaggregation, and reveals its role in sensing coaggregation as a signal. These findings uncover a novel mechanism by which *F. nucleatum* dynamically adapts to polymicrobial environments, offering new perspectives on biofilm formation and bacterial communication in complex oral microbial ecosystems.

## INTRODUCTION

*Fusobacterium nucleatum* is a Gram-negative, spindle-shaped bacterium that plays a central role in oral biofilms, particularly in periodontal disease-associated subgingival plaque, a polymicrobial community of over 400 bacterial species (1–3). It acts as a bridging organism, facilitating the integration of early and late colonizers, a process critical for dental plaque formation (4–6). Beyond its ecological role, *F. nucleatum* is associated with systemic diseases, including preterm birth and colorectal cancer, underscoring its clinical significance (7, 8).

A defining trait of *F. nucleatum* is its ability to coaggregate with diverse bacterial partners through specific surface adhesins, notably RadD and Fap2. Fap2 mediates coaggregation with select species like *Porphyromonas gingivalis* and *Enterococcus faecalis* (9, 10), while RadD enables interactions with a broader range of partners, including early Gram-positive colonizers (*Actinomyces oris*, *Streptococcus sanguinis*, *Streptococcus oralis*, *Streptococcus cristatus*, and *Streptococcus gordonii*), as well as other microbes like *Streptococcus mutans*, *Aggregatibacter actinomycetemcomitans*, and the fungus *Candida albicans* (11–17). RadD also mediates host-pathogen interactions by binding to receptors such as CD147 on colorectal cancer cells (18) and Siglec-7 on natural killer cells (19), highlighting its importance in both biofilm dynamics and pathogenesis.

RadD is encoded by the last gene of the four-gene *radABCD* operon, whose expression is growth-phase-dependent, peaking during the stationary phase (11, 20). Interestingly, deletion of the third gene in this operon, *radC* (also known as *fad-I*), significantly enhances *radD* expression (21). Despite these insights, the regulatory mechanisms governing *radD* expression remain unclear.

Our previous studies identified the CarSR two-component system (TCS) as a regulator of *radD* expression (12). TCSs are critical for bacterial adaptation, consisting of a membrane-bound sensor kinase (SK) and a cytoplasmic response regulator (RR). Upon detecting environmental signals, the SK undergoes autophosphorylation and transfers the phosphate group to its cognate RR, which then modulates target gene expression by binding to promoter regions (22). In *F. nucleatum*, the deletion of *carS* (the SK) represses *radD* expression, while the deletion of *carR* (the RR) enhances it, as revealed by RNA-seq and qRT-PCR analyses (12). These findings suggest that CarS and CarR have opposing roles, with CarS alleviating repression and CarR maintaining it. However, whether CarRS directly regulates *radD*, the broader CarSR regulon, and the environmental signals sensed by this system remain unknown.

To address these gaps, we employed chromatin immunoprecipitation followed by high-throughput DNA sequencing (ChIP-seq) to identify the CarR regulon and determine the consensus DNA sequence recognized by CarR. Our findings demonstrated that CarSR directly regulates *radD* expression and thus influences RadD-mediated fusobacterial coaggregation. Strikingly, we also found that coaggregation itself is sensed by CarSR, creating a feedback loop that further modulates *radD* expression. These discoveries illuminate the molecular mechanisms underlying *F. nucleatum* coaggregation and provide new insights into how this bacterium dynamically adapts to polymicrobial biofilms, enhancing its ecological and pathogenic roles.

## RESULTS

### Construction and Validation of a plasmid-expressing C-terminal FLAG-Tagged CarR

ChIP-seq is a powerful method for studying protein-DNA binding sites in vivo (23, 24). A critical requirement for this technique is a high-quality antibody that specifically recognizes the protein of interest to capture and isolate protein-DNA complexes. Initially, we attempted to generate a CarR-specific polyclonal antibody using purified CarR from *E. coli*. However, this antibody lacked the necessary specificity (Fig. S1). To overcome this limitation, we utilized a FLAG-tagged version of CarR to facilitate detection and interaction studies. FLAG-tag antibodies are commercially available, highly specific, and widely used in ChIP-seq experiments, provided the FLAG tag does not interfere with the fused protein’s function (25, 26).

To this end, we constructed three shuttle plasmids—p*carR*, p*carR-3F*, and p*3F*—using the pCWU6 backbone, each under the control of the native *carSR* promoter (Fig. 1A). These plasmids were designed to express unmodified CarR, CarR with a C-terminal 3xFLAG tag, and the FLAG tag alone, respectively. The constructs were introduced into a *ΔcarR* mutant to evaluate whether the FLAG tag affected CarR’s normal function. FLAG-tagged CarR was explicitly detected by Western blot using a commercially available anti-FLAG antibody (Fig. 1B, middle panel). While the FLAG tag alone was too small (2.4 KDa) to detect via SDS-PAGE, its expression was confirmed through dot blot analysis (Fig. S2). Consistent with previous findings (12), deletion of *carR* significantly increased RadD synthesis (Fig. 1B, lane 2). Complementation with unmodified CarR restored RadD levels to those observed in the wild-type (WT) strain (Fig. 1B, lane 3). Similarly, FLAG-tagged CarR restored RadD levels comparable to those produced by unmodified CarR (Fig. 1B, lane 4). In contrast, the FLAG tag alone did not reverse the *ΔcarR* phenotype (Fig. 1B, last lane), confirming that the FLAG tag itself does not influence RadD production.

**Figure 1:**
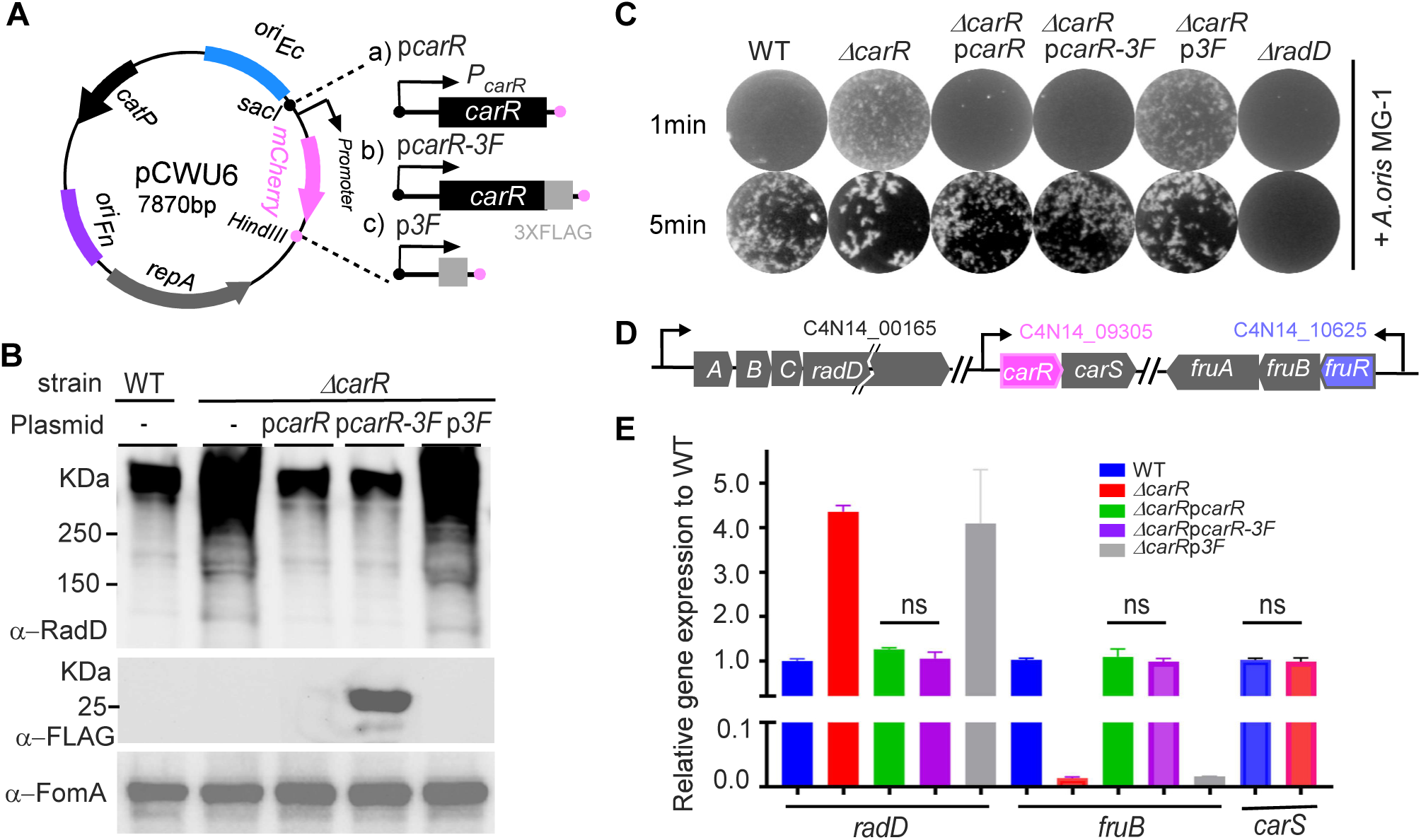
Validation of FLAG-tagged CarR functionality in *F. nucleatum*. **(A)** Schematic of plasmid constructs showing the replacement of *mCherry* in pCWU6 with the gene of interest to generate expression plasmids for wild-type CarR, FLAG-tagged CarR (CarR-FLAG), and FLAG alone, each controlled by the native P*carR* promoter. **(B)** FLAG-tagged CarR mirrors the function of wild-type CarR by restoring RadD expression levels in the Δ*carR* mutant to those observed in wild-type cells. Stationary-phase cultures of the indicated strains (1 mL each) were prepared for SDS-PAGE and Western blotting, using anti-RadD, anti-FLAG, and anti-FomA antibodies. FomA served as the loading control. **(C)** Coaggregation assays show that FLAG-tagged CarR supports coaggregation with *A. oris* MG-1 similarly to untagged CarR. Representative images from three independent experiments demonstrate various coaggregation phenotypes across strains. **(D)** Diagram of the *radABCD*, *carRS*, and *fruRBA* operons, including their relative genomic positions and associated gene IDs (*radD*, *carR*, and *fruB*) as annotated on the NCBI RefSeq assembly: GCF_003019785.1. **(E)** FLAG-tag does not alter CarR’s regulatory effect on the *radD* and *fruRBA* operons at the mRNA level. Cultures of each strain were grown to an OD_600_ of 1.0 in a TSPC medium and then processed for RNA extraction. Transcript levels of *radD* and *fruB* were quantified by qRT-PCR, normalized to *gyrB rRNA*, and set relative to the wild-type, with an arbitrary value of 1. Data represents the mean ± SEM from three biological replicates.

RadD mediates *F. nucleatum* coaggregation with *A. oris*, where higher RadD levels correspond to faster and stronger coaggregation. As shown in Fig. 1C, the *ΔcarR* and *ΔcarR p3F* strains exhibited faster coaggregation than the WT and *carR*-complemented strains. Notably, there were no differences in coaggregation between cells expressing FLAG-tagged and unmodified CarR, both of which mirrored WT behavior.

CarR regulates *radD* expression by controlling the *radABCD* operon (Fig. 1D) at the transcriptional level (12). Previous RNA-seq analysis revealed that CarR regulates over 200 genes, including the *fruRBA* operon involved in fructose utilization (12). Interestingly, CarR exerts opposite regulatory effects on *radD* and *fruRBA*: deletion of *carR* increases *radD* expression but reduces *fruRBA* expression (Fig. 1E) (12). Quantitative RT-PCR confirmed that FLAG-tagged CarR, like unmodified CarR, restored *radD* and *fruB* transcript levels to those observed in WT cells (Fig. 1E). These findings demonstrate that the FLAG tag does not impair fused CarR’s regulatory functions.

### Genome-Wide Identification of the CarR Regulon Using ChIP-seq

With the Δ*carR* strain complemented with FLAG-tagged CarR, we performed ChIP-seq to define the CarR regulon in *F. nucleatum*. Cultures were grown to the late log phase, and formaldehyde was added to crosslink CarR to its DNA binding sites. The cells were collected, lysed, and sonicated to shear the chromosomal DNA into smaller fragments. CarR-DNA complexes were then enriched using anti-FLAG M2 magnetic beads, and the crosslinks were subsequently reversed to purify the DNA. The immunoprecipitated sample (IP) and an unenriched control sample (input) were processed into sequencing libraries. Paired-end sequencing of biological duplicates was performed on the Illumina platform, and the resulting reads were aligned to the *F. nucleatum* subsp. *nucleatum* ATCC 23726 genome. This analysis produced robust and reproducible ChIP-seq peaks (Fig. 2A & S3). A total of 161 significant CarR binding sites (enrichment >2-fold, P < 0.001) were identified across the chromosomes, located in upstream promoter regions (78%, 125 peaks), coding regions (19%, 30 peaks), and overlapping gene end (3%, 6 peaks) (Fig. 2B, Table S3). These results demonstrate that CarR is a global transcriptional regulator in *F. nucleatum*.

**Figure 2:**
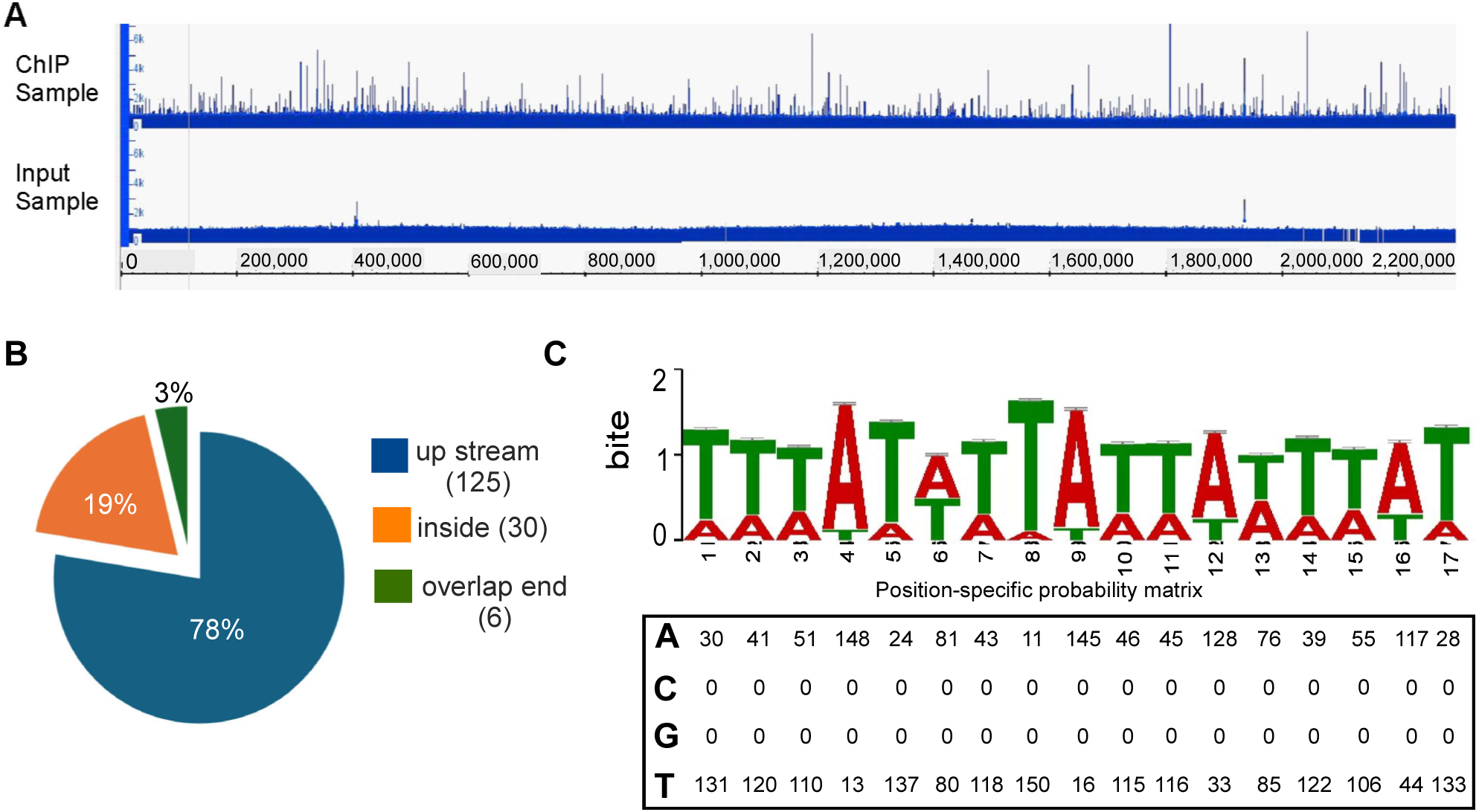
Identification of CarR binding sites in *F. nucleatum* using ChIP-seq. **(A)** A representative genome-wide ChIP-seq profile displaying CarR binding peaks across the F. nucleatum genome is visualized using Integrated Genome Viewer (IGV). Peak height corresponds to the number of reads from the immunoprecipitated (IP) samples (top) and input samples (bottom) mapped to the *F. nucleatum* ATCC 23726 genome, with enriched peaks indicating potential CarR regulatory regions. **(B)** Pie chart representing the distribution of 161 enriched CarR binding peaks, indicating the locations of binding sites within various genes. **(C)** Consensus CarR binding motif generated from ChIP-seq data using MEME. The motif logo shows nucleotide frequency at each position, with letter height representing prevalence. The position frequency matrix illustrates nucleotide distribution across the consensus CarR binding sequence.

Using MEME Suite, we identified a conserved 17-bp AT-rich sequence as the CarR binding motif (Fig. 2C). Previous RNA-seq results revealed that CarR directly or indirectly regulates 236 genes, organized into approximately 80 operons (12). Among these, 40 operons (containing approximately 130 genes) were also identified as CarR targets through ChIP-seq, validating the reliability of the ChIP-seq data (Table 1).

**Table 1.**
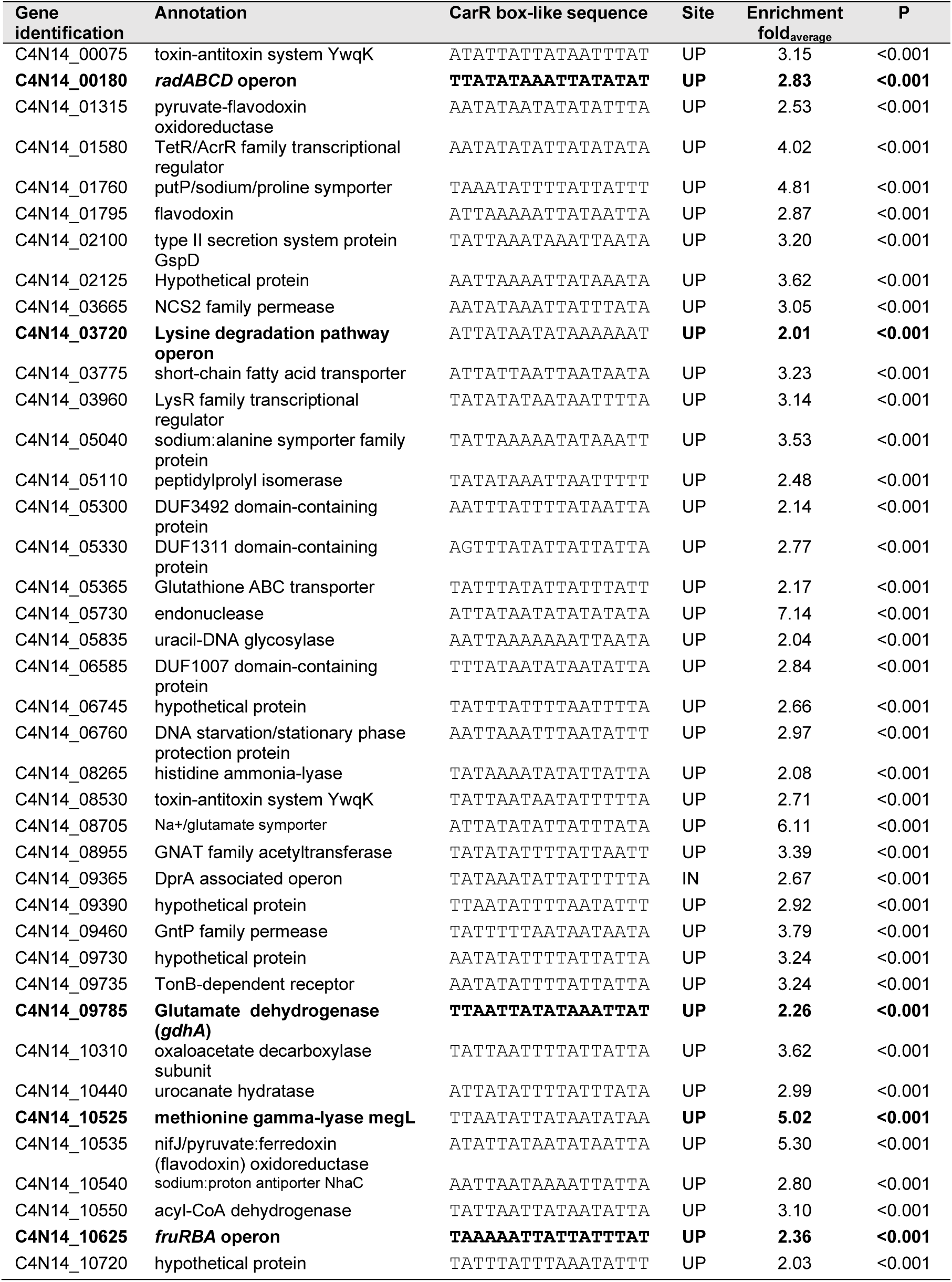
selected. CarR Chip-seq gene list shared with RNA-seq data.

To further validate the ChIP-seq results, we analyzed the expression of several genes detected only by ChIP-seq but not RNA-seq. These included *C4N14_00100* (autotransporter), *C4N14_01790* (glutaredoxin), *clpB*, *C4N14_05075* (*ompA*), *C4N14_05365* (ABC transporter substrate-binding protein), and *C4N14_08250* (response regulator WalR of a WalRK-like TCS). Quantitative RT-PCR confirmed the differential expression of these genes between WT and Δ*carR* strains (Fig. S4). Interestingly, *fap2*, encoded for another major adhesion protein, showed no change in expression, as it was not detected in either RNA-seq or ChIP-seq, confirming that CarR does not regulate it. Collectively, these findings establish CarR as a global transcriptional regulator in *F. nucleatum*, influencing various genes critical for the bacterium’s ecological and functional adaptation in polymicrobial environments.

### CarR Directly Controls RadD Expression

ChIP-seq analysis identified significant enrichment (2.83-fold) at the promoter region of the *radABCD* operon (Fig. 3A), strongly suggesting that CarR binds to this locus. To better characterize the *radABCD* promoter, we first determined its transcription start site using 5’ Rapid Amplification of cDNA Ends (5’ RACE). This start site was located just 8 nucleotides upstream of the ribosome binding site (RBS) of the first gene, *radA*. Sequence analysis also revealed a putative 17-bp CarR binding motif between a predicted −10 box and the transcription start site (Fig. 3B).

**Figure 3:**
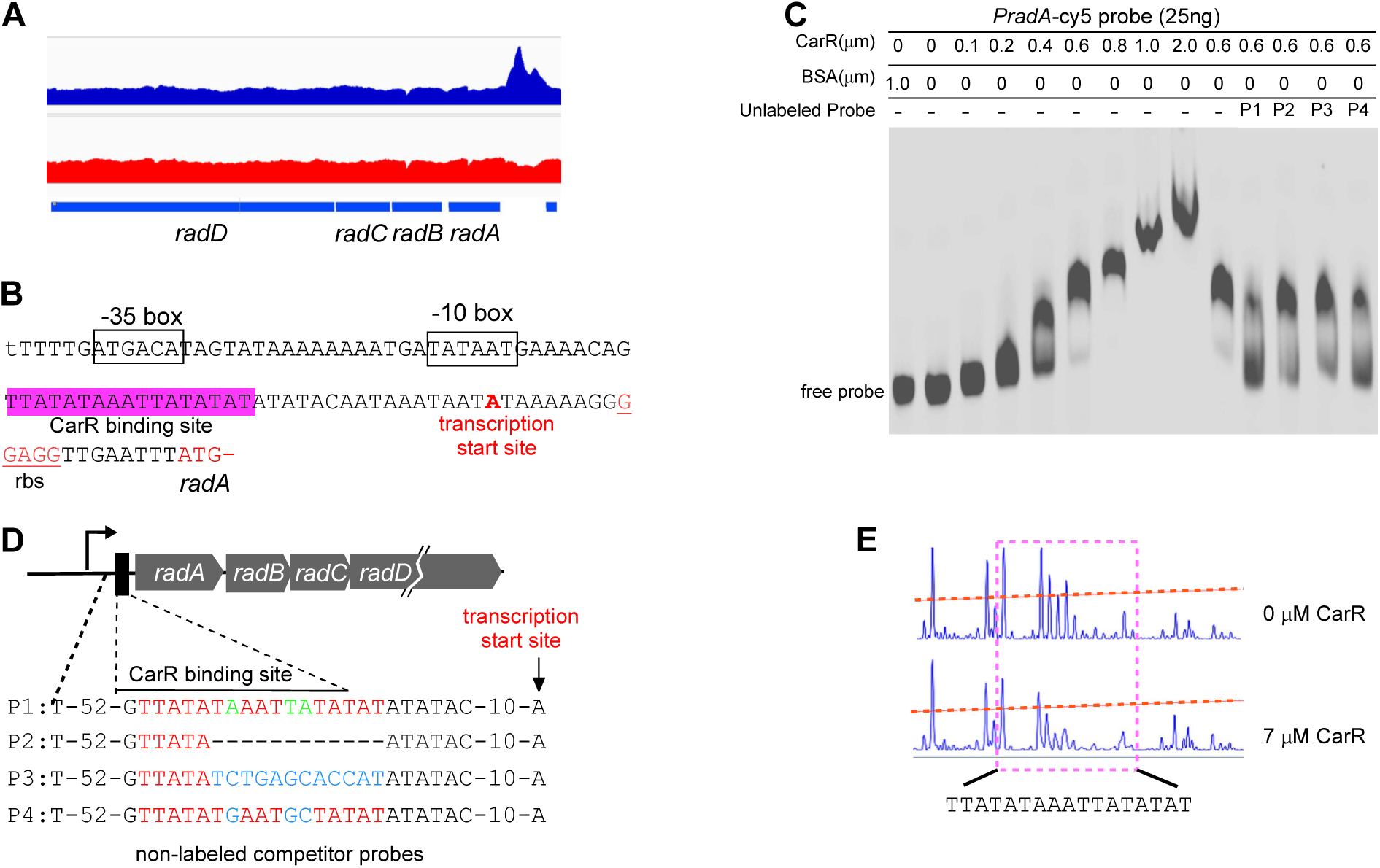
CarR regulation of the *radABCD* operon via direct binding to the *radA* promoter in *F. nucleatum*. **(A)** The ChIP-seq coverage track shows CarR binding enrichment at the *radA* promoter region, visualized using the Integrated Genome Viewer (IGV). The immunoprecipitated (IP) samples (top) display a prominent peak at the *radA* promoter, indicating specific CarR binding. In contrast, the input samples (bottom) serve as controls to confirm the specificity of the enrichment. **(B)** Sequence analysis of the *radA* promoter region, with key regulatory elements annotated. The predicted CarR binding site, highlighted in red, lies between the −10 box and the transcription start site (TSS). Additional elements, including the TSS, ribosome binding site (RBS), start codon of *radA*, and potential −35 and −10 boxes, are also indicated. **(C)** Four synthesized oligonucleotide probes (P1, P2, P3, and P4) were designed for competitive EMSA analysis. P1 represents the wild-type CarR binding site, while P2, P3, and P4 contain distinct mutations within the binding site. **(D)** Electrophoretic mobility shift assay (EMSA) demonstrating CarR binding to the intact *radA* promoter. Purified CarR protein was incubated with a Cy5-labeled *radA* promoter probe in the presence of a 10-fold excess of non-specific competitor DNA (poly[dI-dC]) and 25 ng of labeled probe. Competitive binding was tested by adding a 100-fold excess of unlabeled probes (P1, P2, P3, P4) as indicated, which were pre-incubated with the CarR mixture before the labeled probe. **(E)** DNase I footprinting analysis of CarR binding to the *radA* promoter. Electropherograms display the protected regions of the *radA* promoter following DNase I digestion in the presence 7 µM CarR compared to without CarR, confirming the specific binding region of CarR on the promoter.

To validate CarR’s direct binding to the *radABCD* promoter, full-length CarR protein was produced in *E. coli* and purified to high quality (Fig. S5). Electrophoretic mobility shift assays (EMSAs) were performed using a 215-bp Cy5-labeled DNA fragment spanning the *radABCD* promoter (pr*adA*). Increasing concentrations of CarR resulted in a dose-dependent reduction in the migration rate of the labeled probe, providing clear evidence of binding (Fig. 3C, lanes 2–9). As a negative control, bovine serum albumin (BSA) did not affect probe migration, confirming the specificity of the CarR-DNA interaction (Fig. 3C, lane 1).

To evaluate the functional importance of the 17-bp CarR binding motif, competitive EMSA assays were further conducted using unlabeled competitor probes with specific mutations (P1– P4; Fig. 3D). P1, containing the wild-type binding sequence, effectively outcompeted the labeled probe for CarR binding, reversing the migration shift. In contrast, P2, lacking 12 bp of the motif, and P3, which included 9 nucleotide mutations, exhibited little competition. P4 reversed most of the shift, containing mutations in the 3 most conserved nucleotides (Fig. 2C). These findings demonstrate that the 17-bp motif is essential for CarR binding, with the conserved nucleotides playing a critical role in maintaining the interaction.

To pinpoint the exact CarR binding site, a dye/primer sequencing-based DNase I footprinting assay was further performed using the same p*radA* DNA probe employed in the EMSA studies, this time labeled with carboxyfluorescein [6-FAM]). Comparative analysis of the electropherograms, with and without CarR, revealed that CarR protects the 17-bp motif from DNase I digestion, confirming it as the precise binding site (Fig. 3E). Together, these findings provide compelling evidence that CarR directly regulates *radABCD* expression by binding specifically to CarR-binding motif in its promoter region, highlighting its role as a direct transcriptional regulator of *radD*.

### CarR Direct Regulation of Operons Involved in Fructose and Amino Acid Utilization

To further validate the ChIP-seq findings and explore the dual regulatory roles of CarR— repressing genes such as *radD* while activating others like *fruRBA*—we examined the promoter region of the *fruRBA* operon using EMSAs (Fig. 4A). The *fruRBA* operon widely presented in oral bacteria (27, 28), consists of three genes: *fruR* (encoding a transcriptional regulator), *fruB* (encoding a component of the phosphotransferase system (PTS) specific to fructose), and *fruA* (encoding the fructose-specific permease of PTS). These genes enable *F. nucleatum* to utilize fructose (29). While *F. nucleatum* prefers to utilize amino acids such as glutamate and lysine for ATP production and growth (1), it selectively uses fructose as its only sugar-based energy source (Wu et al., unpublished data). Supplementing the growth medium with fructose—but not glucose—significantly enhances the growth of *F. nucleatum* (Fig. 4B, column 1 vs 6). Deletion of *fruB* abolished the bacterium’s ability to use fructose. In contrast, complementation restored this function (Fig. 4B, column 4 vs. 5 and column 10 vs. 11). Consistent with previous observations that CarR positively regulates *fruRBA* expression (Fig. 1D), the Δ*carR* mutant was also unable to utilize fructose (Fig. 4B, columns 6 and 12).

**Figure 4:**
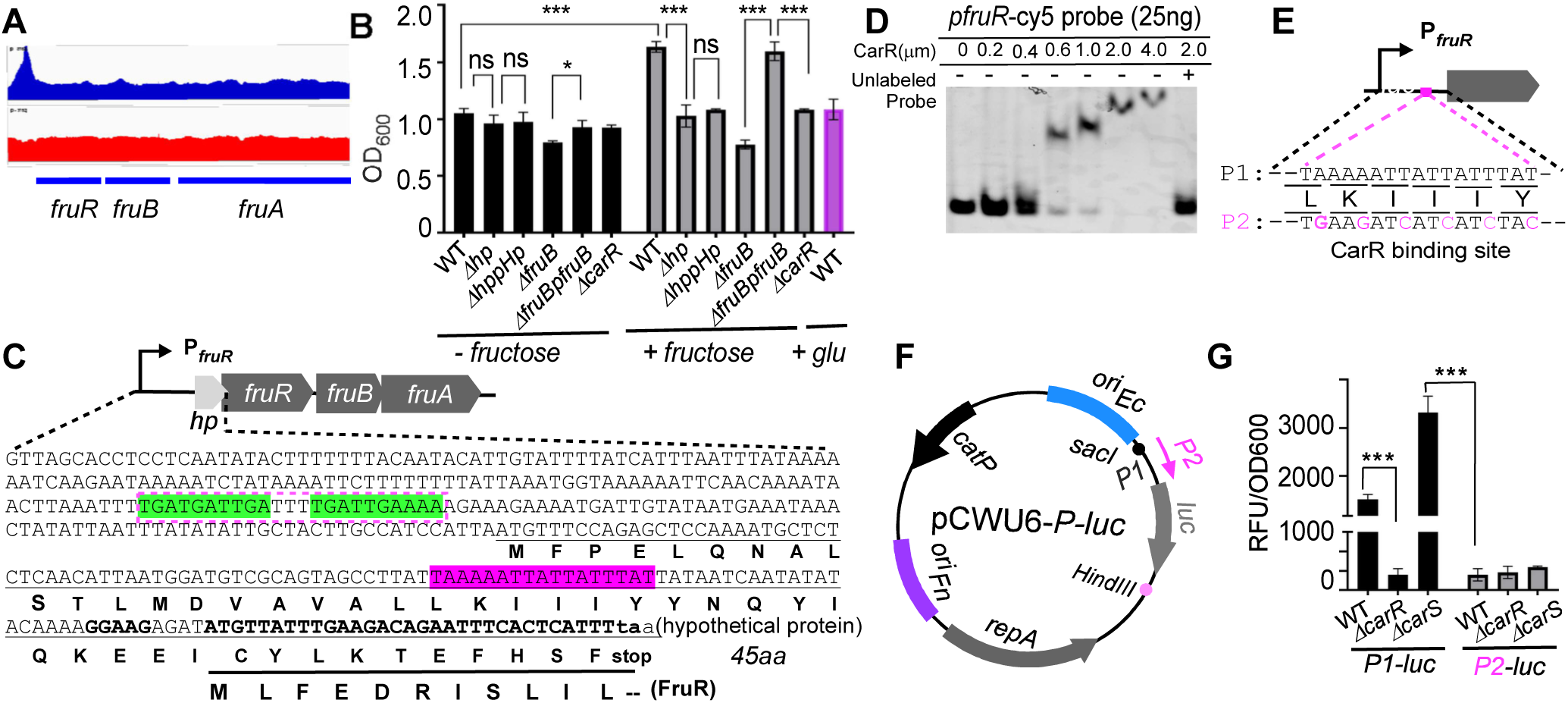
CarR regulation of the *fruRBA* operon via direct binding to the *fruR* promoter in *F. nucleatum*. (A) ChIP-seq analysis shows CarR binding at the *fruR* promoter. The ChIP samples (top) display a clear peak at the promoter, indicating CarR binding, while the input samples (bottom track) confirm this enrichment is specific. Data from the two experiments are consistent, as shown in the Integrated Genome Viewer (IGV). (**B)** Growth assay demonstrating that deletion of the hypothetical protein region upstream of the *fruRBA* operon abolishes fructose-stimulated growth in *F. nucleatum*. This phenotype cannot be restored by complementation, suggesting possible misannotation of this region. Growth was measured as OD*600* after 12 hours in the presence of 20 mM fructose or TSPC alone. **(C)** Sequence analysis of the *fruRBA* promoter region with annotated regulatory elements. The predicted CarR binding site, shaded in red, is located within a region encoding a hypothetical 45-amino-acid protein. This hypothetical protein’s coding sequence overlaps with the *fruR* gene. Potential FruR binding sites are highlighted in green**. (D)** EMSA demonstrating CarR binding to the intact *fruR* promoter. Purified CarR protein was incubated with a Cy5-labeled *fruR* promoter probe in the presence of a 10-fold excess of non-specific competitor DNA (poly[dI-dC]) and 25 ng of the labeled probe. As indicated, competitive binding was assessed by pre-incubating the CarR mixture with a 100-fold excess of unlabeled probes before adding the labeled probes. (**E)** Schematic of two *PfruR* promoter versions: wild-type and a variant with a mutated CarR binding site, both used to drive luciferase expression as shown in panel F. P1 represents the wild-type CarR binding site within *fruR* promoter, while P2 contains distinct mutations within the binding site. **(F)** Schematic of plasmid constructs for luciferase reporter assays, showing the replacement of *mCherry* in pCWU6 with luciferase, driven by the wild-type or mutated *fruR* promoter. **(G)** Promoter activity of *PfruR* and its mutant variant fused to luciferase in WT, Δ*carR*, and Δ*carS* backgrounds. Cultures were sampled at an OD_600_ of 0.6 for luciferase assays. Luciferase activity (RLU) was normalized to cell density (OD600), with data representing the mean ± SD from three biological replicates.

ChIP-seq analysis revealed a 2.36-fold enrichment in the upstream region of *fruR* (Fig. 4A and Table 1). Interestingly, this region was annotated in the BioCyc database (www.biocyc.org) as encoding a hypothetical 45-amino-acid (aa) protein (HP), overlapping with the first 11 amino acids of FruR (Fig. 4C). Notably, a predicted CarR-binding site (highlighted in red in Fig. 4C) is located within this region.

To determine whether the *hp* gene is functional, we examined its potential for translation. Typically, a functional gene has a ribosome binding site (RBS) near its start codon. However, we could not identify such an RBS upstream of the *hp* start codon. We deleted the region encoding the HP protein’s 2nd to 23rd amino acids to investigate further, ensuring that the *fruR* RBS remained unaffected. This deletion significantly impaired fructose utilization, yielding a phenotype similar, though not identical, to that of the *fruB* mutant (Fig. 4B). Unlike the *fruB* mutant, which completely abolishes fructose utilization, the deletion of *hp* retains a residual ability to utilize fructose. However, the ectopic expression of the intact *hp* gene under its native promoter failed to restore normal fructose utilization (Fig. 4B). These findings suggest that the region containing CarR-binding motif upstream of *fruR* likely serves as a regulatory element for *fruRBA* expression rather than encoding a functional protein.

To confirm that CarR directly binds the upstream region of *fruR*, we PCR amplified a 131-bp DNA fragment containing the 17-bp CarR-binding motif and labeled it with Cy5. The labeled probe was then incubated with increasing concentrations of CarR protein for EMSA assays. The results showed a dose-dependent shift in probe migration, indicating CarR binding (Fig. 4D, lanes 1-7). Furthermore, excess unlabeled probes successfully competed with the labeled probes, preventing the shift and confirming the specificity of CarR binding to the *fruRBA* promoter (Fig. 4D, the last lane).

To assess the importance of the CarR-binding motif, we constructed two versions of the *fruRBA* promoter (a 330-bp fragment upstream of the *fruR* start codon): a wild-type version (P1) and a mutated version (P2) containing a 6-bp alteration in the CarR-binding motif (Fig. 4E). The mutations in P2 were designed to be silent for the *hp* gene, ensuring that any observed effects reflected regulatory changes rather than disruption of a potential coding sequence—despite evidence suggesting that *hp* is likely misannotated. Both promoter fragments were fused to a luciferase reporter gene and subcloned into pCWU6, generating plasmids pCWU6-*P1-luc* and pCWU6-*P2-luc* (Fig. 4F), which were introduced into WT, Δ*carR*, and Δ*carS* strains.

Luciferase activity from P1 was significantly reduced in the Δ*carR* mutant compared to WT, confirming that CarR positively regulates the *fruRBA* promoter (Fig. 1E and 4G). In contrast, luciferase activity was dramatically increased in the Δ*carS* mutant, suggesting that CarS represses *fruRBA* transcription. These results align with our qRT-PCR results (Fig. 1E & S6). However, these regulatory effects were absent in strains containing the P2 promoter, where luciferase activity remained consistently low across all backgrounds, including WT. Notably, P2 activity in WT cells was comparable to P1 activity in the Δ*carR* mutant, emphasizing the essential role of the carR-binding site for promoter activation.

To further reinforce the validity of our ChIP-seq data and expand the evidence supporting CarR binding to the identified CarR-binding motif-containing promoters, we performed EMSAs using purified CarR protein on three selected promoters: *megL*, *gluD*, and *ldp* (lysine degradation pathway). The *megL* gene encodes methionine gamma-lyase, known to play a role in cysteine metabolism in *F. nucleatum* (30). *gdhA* encodes glutamate dehydrogenase, involved in glutamine utilization, and *ldp* is crucial for lysine metabolism in this bacterium (31). ChIP-seq analysis revealed significant enrichment in the promoter regions of these operons, each containing a 17-bp CarR-binding motif (Fig. 5A, Tabe 1). EMSAs confirmed that CarR specifically binds to the promoter regions of *megL*, *gdhA*, and *ldp*, as evidenced by evident probe retardation (Fig. 3B-D). In contrast, no binding was observed with the *fap2* promoter, which was used as a negative control since CarR does not regulate *fap2* expression (Fig. 5E). These results confirm that CarR directly regulates these operons involved in fructose and amino acid utilization, including cysteine, glutamine, and lysine metabolism pathways.

**Figure 5.**
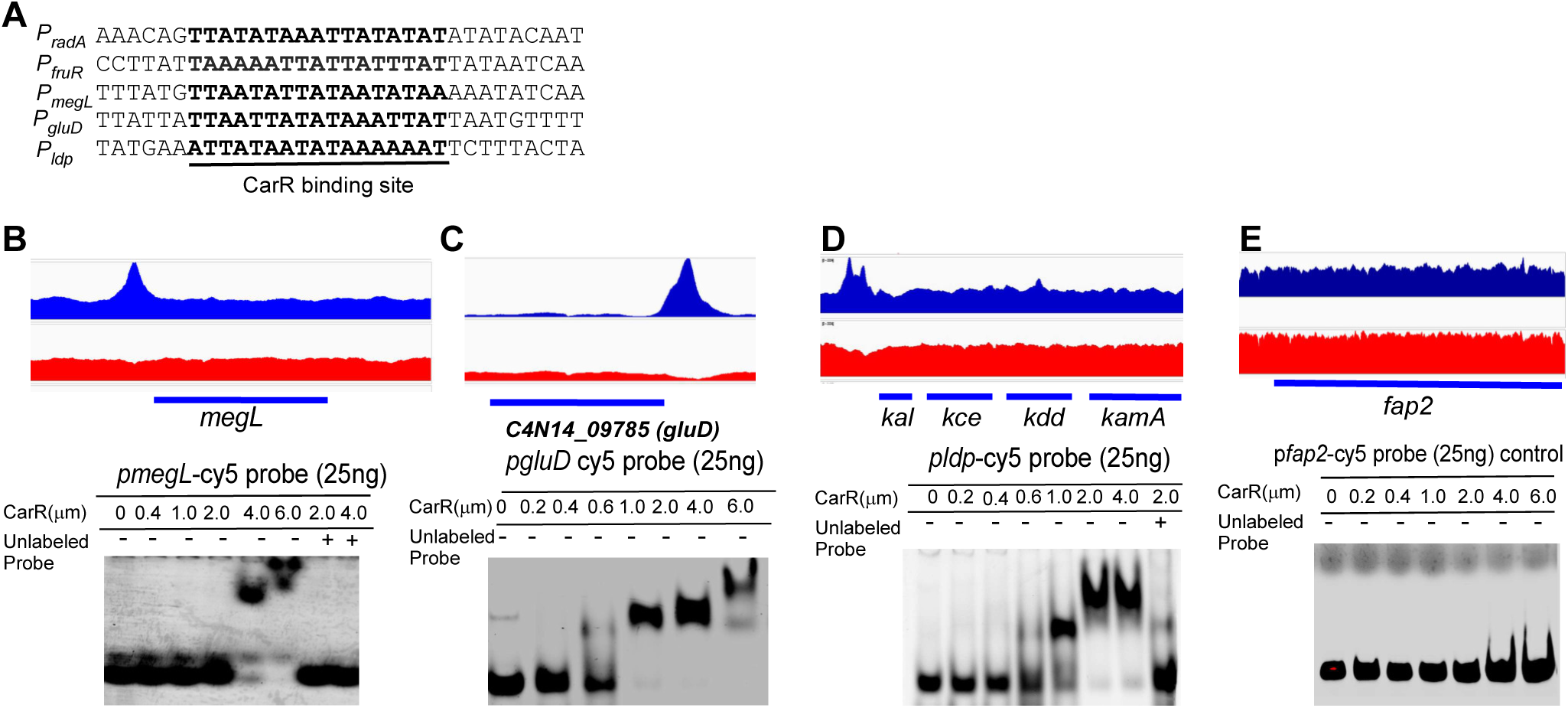
Validation of ChIP-seq findings by EMSA, confirming CarR’s regulatory role in *F. nucleatum* metabolic pathways through direct promoter binding. (A) Alignment of ChIP-enriched regions from five target genes identifies a conserved CarR binding motif, underlined and highlighted in black, demonstrating sequence specificity in CarR binding. **(B, C, D, E)** Combined ChIP-seq and EMSA analyses reveal CarR binding at the *megL*, *gluD*, and *kal* promoters, while showing no binding at the *fap2* promoter, which is used as a negative control. Top panels: ChIP-seq coverage tracks from two independent experiments (ChIP sample in blue and input sample in red) exhibit consistent enrichment at the promoter regions of *megL*, *gluD*, and *kal*, mapped to the *F. nucleatum* ATCC 23726 genome. Data are visualized using the Integrated Genome Viewer (IGV). Bottom panels: EMSA results confirm CarR binding to Cy5-labeled promoter probes (*megL*, *gluD*, and *kal*) in a concentration-dependent manner. A 10-fold excess of non-specific competitor DNA was included to validate specificity, and competitive binding was further confirmed by pre-incubation with a 100-fold excess of unlabeled probes. The *fap2* promoter is a negative control, demonstrating no interaction with CarR.

### Coaggregation Enhances *radD* Expression via the CarSR Two-Component System

Bacterial TCSs sense environmental signals through sensor kinases’ signal-sensing domain (SSD) (22). In *F. nucleatum*, the sensor kinase CarS contains two transmembrane (TM) domains at its N-terminus. The region between the two TM domains, approximately 106 amino acids long, is predicted to be periplasmic and serves as the SSD (12). A BLAST search using the CarS SSD sequence revealed that this sequence is unique to *F. nucleatum* and not associated with any known functions in other TCS systems. Given that the CarSR system regulates *radD* expression and RadD is essential for coaggregation between *F. nucleatum* and many Gram-positive oral bacteria, we hypothesized that CarS senses RadD-mediated coaggregation and transduces this signal to CarR.

To test this hypothesis, *F. nucleatum* was cultured alone or co-cultured with *Actinomyces oris* MG-1 in a TSPC medium (32). Co-cultures were prepared in three configurations: (1) simple mixing, (2) forming a high-density cell pellet, and (3) generating visible coaggregation clumps (Fig. 6B). After 2 hours of incubation, RNA was extracted for qRT-PCR analysis of *radB* expression, which serves as a proxy for *radD* expression due to the experimental use of *radD*-engineered strains (Fig. 6A). This approach ensures a unified analysis of the data, allowing consistent interpretation across experimental conditions.

**Figure 6:**
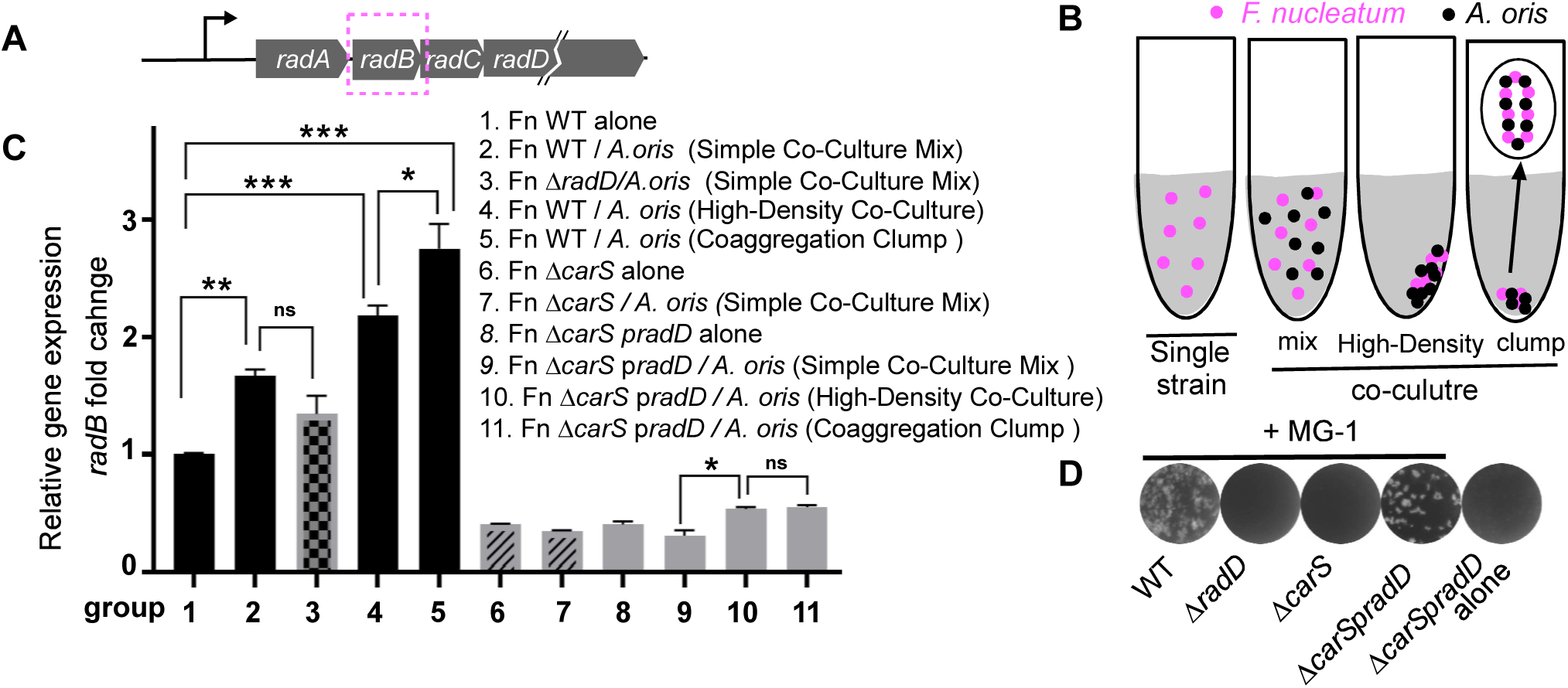
Coaggregation as a signal sensed by the CarRS two-component system in *F. nucleatum*. **(A)** Diagram of the *radABCD* operon structure, with the *radB* gene highlighted in a box to indicate the target detection region for qRT-PCR. **(B)** Schematic of the experimental design, illustrating *F. nucleatum* single-strain and three co-culture conditions with *A. oris* MG-1 used to investigate coaggregation signaling. Co-culture conditions include: (1) Simple Co-Culture Mix, (2) High-Density Co-Culture, and (3) Coaggregation Clump Formation. Detailed descriptions of each condition are provided in the Methods section. **(C)** qRT-PCR analysis of *radABCD* operon expression across the different experimental conditions, with RNA extracted from each indicated experimental setup. *radB* expression levels were normalized to the *gyrB* gene, and the experiments were validated in two independent runs, each conducted in triplicate. The results are shown as means ±SD (n=3). *, P<005, **, P<0.01, n.s, not significant (student’ *t* test) **(D)** Coaggregation assay showing that RadD expression in the Δ*carS* mutant restores coaggregation ability to levels similar to those of the wild-type strain in coaggregation with *A. oris* MG-1. Representative images from three independent experiments with the indicated strains.

When *F. nucleatum* WT was mixed with *A. oris*, *radB* expression significantly increased compared to WT cultured alone (Fig. 6C, columns 2 vs. 1). However, since fresh TSPC medium contains lysine or lysine-containing peptides that interfere with coaggregation (12), this increase in *radB* expression may not be explained due to coaggregation. Supporting this, the simple mix of *F. nucleatum* Δ*radD* (coaggregation-defective mutant) with *A. oris* also led to increased *radB* expression, albeit slightly lower than in the WT co-culture, with no statistical significance (columns 2 vs. 3). To confirm that coaggregation enhances *radD* expression unequivocally, we created a coaggregation-state co-culture. WT *F. nucleatum* and *A. oris* were coaggregate in a coaggregation buffer to form visible clumps. The buffer was then carefully replaced with fresh TSPC medium without disrupting the clumps, and the culture was incubated for another 2 hours. This coaggregation-state culture displayed significantly higher *radB* expression than both WT alone and the simple mix co-culture (Fig. 6C, columns 5 vs. 1 and 2). Since coaggregation inherently increases cell density, which could alter gene expression, we tested whether high cell density alone could explain this effect. High-cell-density cultures were prepared by centrifuging cells from a simple mix. While high cell density moderately enhanced *radB* expression, the effect was significantly less pronounced than in coaggregation-state cultures (columns 4 vs. 5). These results suggest that RadD-mediated coaggregation specifically enhances *radD* expression.

To determine if this phenomenon is specific to *A. oris*, we repeated the experiments using another oral bacterium, *S. oralis*. Similar results were observed, with *radD* expression significantly increasing during coaggregation with *S. oralis* (Fig. S7). This suggests that coaggregation-dependent *radD* expression may be a universal feature of *F. nucleatum* in interactions with Gram-positive partners.

We next investigated whether CarS is required for this coaggregation-dependent regulation. Δ*carS* mutants, which produce reduced levels of RadD (31), failed to form coaggregation-state cultures with *A. oris* (Fig. 6D). To restore coaggregation, we constructed a Δ*carS* strain with ectopic *radD* expression driven by a constitutive promoter (Δ*carS* p*radD*). This strain restored coaggregation to WT levels (Fig. 6D). Consistent with the reduced RadD levels in Δ*carS*, *radB* expression was significantly lower in Δ*carS* cells compared to WT cells cultured alone (Fig. 6C, column 6 vs. 1). Simple mixing of Δ*carS* with *A. oris* did not increase *radB* expression (columns 6 vs. 7). Although coaggregation-state cultures of Δ*carS* p*radD* with *A. oris* modestly increased *radB* expression compared to simple mixes, the increase was much lower than in WT coaggregation-state cultures (columns 5 vs. 10). No difference was observed in high-cell-density cultures (columns 10 vs. 11). These findings strongly support that RadD-mediated coaggregation enhances *radD* expression through the CarSR system.

## DISCUSSION

*F. nucleatum* plays a pivotal role in oral biofilm formation, with RadD-mediated coaggregation being essential to this process. Our prior RNA-seq analysis indicated that the CarSR TCS regulates *radD* expression. However, RNA-seq alone cannot discern whether this regulation is direct or indirect. In this study, we employed ChIP-seq to elucidate the CarR regulon and confirm the direct regulation of *radD* by CarR. Additionally, we discovered that CarS senses RadD-mediated coaggregation, which subsequently enhances *radD* transcription.

### CarR as a Global Regulator

Studying a bacterial TCS needs to answer two key questions: which genes are in its regulon, and what signals it detects. To determine the CarR regulon and assess direct control over *radD* expression, we constructed a strain expressing a functional FLAG-tagged CarR. Then we used it to perform ChIP-seq analyses (Fig. 1). This approach identified 161 gene operons, encompassing over 400 genes constituting approximately 20% of the *F. nucleatum* ATCC 23726 genome (totaling about 2,180 genes) (Table 1 and S3). These targeted operons are distributed across the chromosome (Fig. 2A) and involve diverse functions, including genetic and environmental information processing, cellular processes, and metabolism (Table S3). This extensive involvement indicates that CarR is a global regulator in *F. nucleatum*.

Notably, 40 gene operons (approximately 130 genes) identified by ChIP-seq were also detected in our previous RNA-seq analysis, which found that CarR controls about 236 genes (31). This overlap suggests that nearly half of the genes regulated by CarR are indirectly controlled, potentially through other DNA regulators directly regulated by CarR.

### CarR Directly Regulates *radD* Expression

Through MEME suite analysis, we identified a conserved 17-bp AT-rich CarR-binding motif located in the promoter region of the *radABCD* operon, positioned between a predicted −10 box (TATAAT) and the transcriptional start site (TSS) (Fig.3B). The unusually long distance (41 bp) between the −10 box and TSS deviates from the typical 5-9 bp found in bacterial promoters (33). This discrepancy raises the possibility that our predicted −10 box location is incorrect or that the motif itself overlaps or includes the −10 box, as both sequences are AT-rich. Further experimental validation is required to clarify this architecture.

Nonetheless, electrophoretic mobility shift assays (EMSA), mutagenesis, and DNase I footprinting assays confirmed that this 17-bp motif in the *radABCD* promoter is crucial for CarR binding (Fig.3), indicating that CarR directly regulates *radD* expression.

### Opposing Effects of CarS and CarR Deletions

Deletion of *carR* results in increased *radD* expression, whereas deletion of *carS* leads to decreased *radD* expression (31). Typically, deleting either the sensor kinase or the response regulator of a TCS yields similar phenotypes, as both components are part of the same signaling pathway (22). The opposing effects observed here are intriguing and unexplained, necessitating further investigation. In *S. mutans*, the CiaRH TCS exhibits opposing effects on mutacin production and competence development when either component is mutated because CiaH represses CiaR expression; deletion of *ciaH* increases CiaR expression, leading to opposite phenotypes (34). However, in our study, deletion of *carS* or *carR* did not affect each other’s expression (Fig 1E & S7), suggesting a different mechanism. One possibility is crosstalk between different TCSs, which requires further exploration (35).

Based on our current data, we propose a model where non-phosphorylated CarR acts as a repressor of *radABCD* expression while phosphorylated (activated) CarR serves as an activator (Fig. 7A). The ratio between these two states may determine the level of *radABCD* expression. This hypothesis posits that both non-activated and activated CarR bind to the CarR-binding motif but induce different DNA conformational changes, leading to repression or activation. Ongoing experiments aim to validate this model.

**Figure 7.**
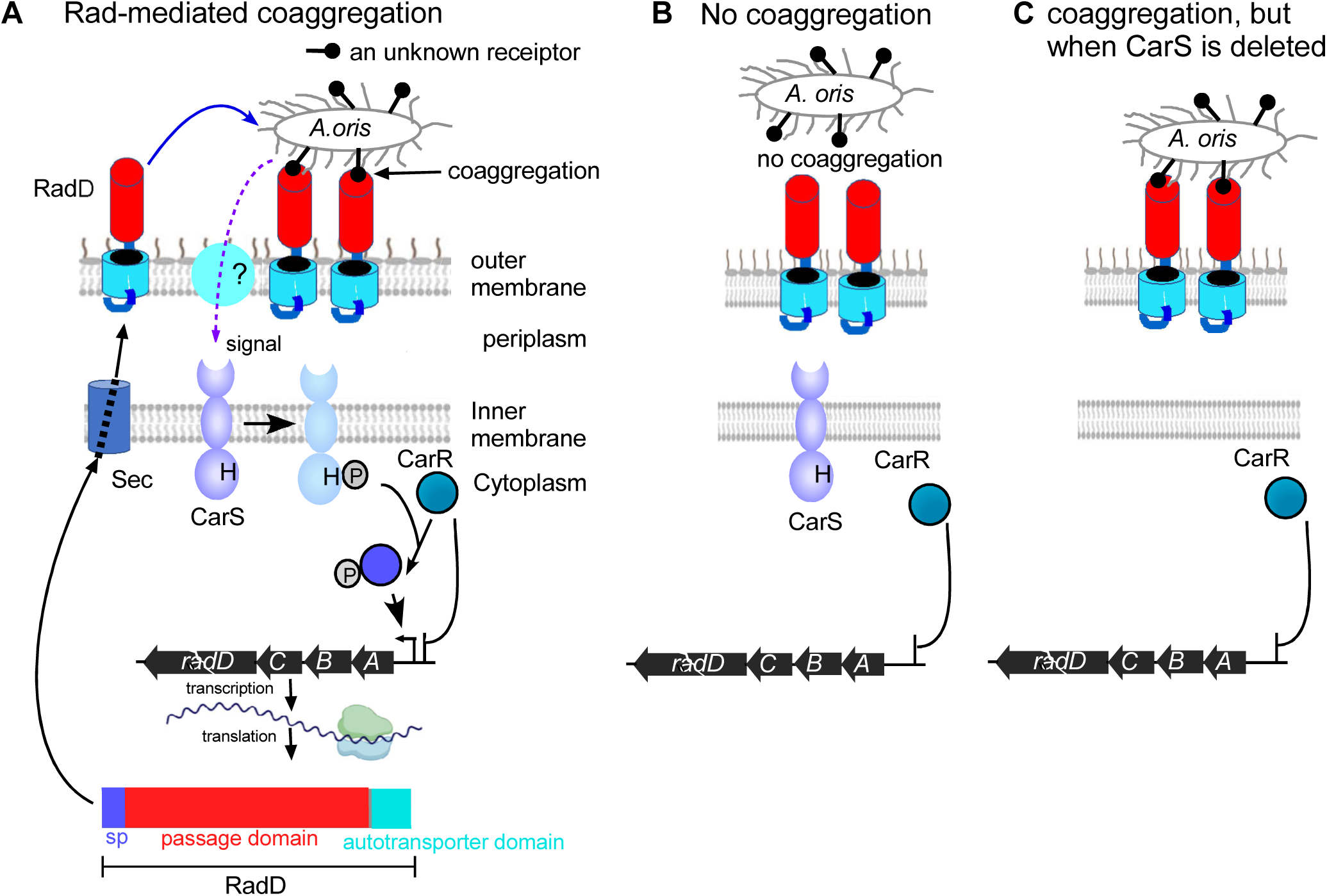
Model of CarS-mediated sensing of RadD-dependent coaggregation and regulation of *radD* expression. **(A)** In wild-type *F. nucleatum*, RadD interacts with an unknown receptor on *A. oris*, initiating coaggregation. This coaggregation generates a signal that activates CarS, leading to CarR-mediated transcriptional regulation of *radD*. RadD expression forms a positive feedback loop, enhancing coaggregation. See main text for details. **(B)** In the absence of coaggregation, CarS remains inactive, leaving CarR in its non-phosphorylated state. Non-phosphorylated CarR is a repressor of the *radABCD* operon, leading to reduced *radD* expression. **(C)** Coaggregation still occurs without CarS (Δ*carS* p*radD*), but the signal cannot be relayed to CarR. As a result, CarR remains non-phosphorylated and unable to activate *radD* transcription, leading to repression of *radD* expression despite coaggregation.

### Dual Regulation of *radD* and *fruRBA* by CarR

The deletion of *carR* results in increased *radD* expression but decreased *fruRBA* expression (Fig. 1E), uncovering a fascinating regulatory dynamic. While *fruRBA* is directly controlled by CarR and enables *F. nucleatum* to utilize fructose for growth, the bacterium exhibits a metabolic preference for amino acids such as glutamine and lysine (1). Notably, operons involved in glutamate and lysine utilization are also directly regulated by CarR (Fig. 5), but their expression follows a pattern opposite to that of *fruRBA* (12).

Given *radD*’s essential role in mediating coaggregation with bacterial partners such as *A. oris*, *S. oralis*, and *S. gordonii*, an ecological rationale for this regulation emerges. These coaggregation partners could also use fructose for growth through their respective *fruRBA* operons (36). By downregulating its own fructose utilization upon coaggregation via RadD, *F. nucleatum* might strategically minimize competition for this nutrient. Instead, it prioritizes amino acid metabolism, thereby facilitating its partners’ fructose consumption and promoting the overall growth of the polymicrobial community in the oral cavity.

This cooperative metabolic adjustment underscores the potential ecological significance of CarR’s dual regulation of *radD* and *fruRBA*. We hypothesize that this regulatory interplay reflects an adaptive strategy that enables *F. nucleatum* to balance nutrient utilization with fostering community dynamics in oral biofilms. Future studies will dissect the molecular mechanisms underlying CarR’s regulation of *fruRBA*, *radD*, *gluD*, and *ldp* to further elucidate their contributions to the ecological roles and success of *F. nucleatum* within polymicrobial environments.

### CarS Senses Coaggregation

The signal-sensing domain (SSD) of CarS in *F. nucleatum* is unique, lacking homology to known SSDs in other bacterial two-component systems (TCS). This distinctiveness suggests that CarS may detect specific signals pertinent to *F. nucleatum*’s ecological niche. Notably, the involvement of a TCS in bacterial coaggregation has not been previously reported. To investigate whether RadD-mediated coaggregation serves as a signal sensed by CarS, we conducted a series of experiments.

When wild-type *F. nucleatum* was co-cultured with *A. oris* in a simple mixture, an increase in *radB* expression—a proxy for *radD*—was observed (Fig. 6, lane 2). However, this upregulation was not attributed to RadD-mediated coaggregation, as co-cultures with a coaggregation-defective mutant also showed elevated *radB* levels (Fig. 6, lane 3). These findings imply that the observed upregulation may result from increased cell density, aligning with previous observations that RadD expression peaks during the stationary phase when cell density is highest (20). Supporting this, high-density cell pellet co-cultures exhibited significant *radB* upregulation (Fig. 6, lane 4), suggesting a possible quorum sensing mechanism linked to cell density that may also regulate RadD expression. However, the specific signal remains unidentified, particularly as this *F. nucleatum* strain does not produce AI-2 (37) and has not been shown to produce acyl homoserine lactones (AHLs) (38).

Interestingly, these results contradict prior studies where co-culturing *F. nucleatum* with *S. gordonii* led to reduced *radD* expression (21). Such discrepancies may stem from differences in co-culture partners, media conditions, or incubation times, necessitating further investigation.

To directly assess whether RadD-mediated coaggregation enhances *radD* expression, we established coaggregation-state cultures using WT *F. nucleatum* or Δ*carS* p*radD* strains with *A. oris* (Fig. 6). Our data confirm that RadD-mediated coaggregation significantly elevates *radB* expression and that this enhancement depends on CarS. Based on these findings, we propose the following model (Fig. 7A): Signal detection begins when WT *F. nucleatum* aggregates with *A. oris* through interactions between RadD and an unidentified receptor on *A. oris*. This coaggregation generates a signal that is transmitted either directly to the SSD of CarS or via an intermediary molecule. Upon signal detection, CarS undergoes autophosphorylation, activating itself. The activated CarS subsequently transfers the phosphate group to CarR, which binds to the CarR-binding motif in the *radABCD* promoter to initiate operon transcription. RadD is translated, secreted via the Sec machinery, and anchored in the outer membrane. Increased RadD levels further enhance coaggregation, creating a positive feedback loop that amplifies *radD* expression. This mechanism strengthens physical interactions between *F. nucleatum* and its partners, promoting polymicrobial biofilm formation. Without coaggregation, CarS remains inactive, leaving CarR in its non-phosphorylated form, which represses *radABCD* expression (Fig.7B). Similarly, if coaggregation occurs without CarS, the signal does not reach CarR, keeping it inactive and *radD* expression repressed (Fig. 7C).

While we have demonstrated this phenomenon with *A. oris* and *S. oralis*, it remains unclear whether this mechanism applies universally to other RadD-dependent coaggregation partners. In other words, are all RadD-mediated bacterial coaggregation events interpreted as the same signal? If not, how does *F. nucleatum* distinguish between different coaggregation partners? An even more intriguing question is whether Fap2-mediated coaggregation can also be sensed through CarS to enhance *radD* expression. If it cannot, how does *F. nucleatum* differentiate between coaggregation mediated by these two distinct adhesins? Additionally, recent studies have shown that *F. nucleatum* uses RadD to interact with host cells (18, 19). Could RadD-mediated host interactions be perceived as a type of coaggregation, and might these signals be sensed by CarS to upregulate *radD* expression? These compelling questions highlight important directions for future research.

In summary, our findings reveal that the CarSR TCS directly regulates *radD* expression and that RadD-mediated coaggregation serves as a signal sensed by CarS to enhance transcription. This study provides new insights into the molecular mechanisms underlying *F. nucleatum*’s role in polymicrobial biofilms and its interactions with microbial and host partners.

## MATERIALS AND METHODS

### Bacterial strains, media, and growth conditions

Bacterial strains and plasmids used in this study are listed in Table 1. *F. nucleatum* strains were grown in tryptic soy broth (TSB) supplemented with 1% Bacto peptone plus 0.25% freshly made cysteine (TSPC) or on TSPC agar plates in an anaerobic chamber filled with 80% N_2_, 10% H_2_, and 10% CO_2_. *A. oris* MG-1 was grown in heart infusion broth (HIB). *S. oralis* #34 and *S. gordonii* DL1 were grown in the Brain Heart Infusion medium. *E. coli* strains were grown in Luria-Bertani (LB) broth. Antibiotics used as needed were chloramphenicol (15 μg mL^−1^) and thiamphenicol (5 μg mL^−1^). Reagents were purchased from Sigma unless indicated otherwise.

### Plasmid construction

All plasmids used in this study were constructed using Gibson assembly cloning, following the manufacturer’s protocol with NEBuilder HiFi DNA Assembly Master Mix (E2621L). This method requires adjacent DNA segments to share at least 18-bp of identical sequences at their ends. These sequences were introduced through PCR amplification, using primers with 5′ ends complementary to adjacent segments and 3′ ends that anneal to the gene-of-interest sequence. PCR reactions were carried out with a 2X Hot Start PCR Master Mix (TaKaRa; catalog no. R405A). A list of primers used for PCR is provided in Table S2. The Gibson assembly reactions were incubated at 50°C for 20–60 minutes, after which 5 µL of the assembled product was used to transform *E. coli* DH5α competent cells. The authenticity of each resultant plasmid was confirmed by DNA sequencing, and validated plasmids were subsequently transferred to fusobacterial strains via electroporation (39). All oligonucleotide primers (Table S2) were custom-synthesized by Sigma Aldrich.

i. p*carR*, pcar*R-3F* and *p3F*. Primers pcarR3xF-F and pcarR-R amplified a fragment (P-carR) containing the carR promoter and its ORF (C4N14_09305, at www.biocyc.org). Using primers pcarR3xF and carR-3xF-R1, another fragment was amplified with a 3xFLAG tag appended to the CarR C-terminus (P-carR::3xFLAG). P-carR and P-carR::3xFLAG were cloned into KpnI/XhoI-digested pCWU6 via Gibson assembly to yield plasmids p*carR* and p*carR-3F*, respectively. For the control plasmid p*3F*, inverse PCR with primers PcarR-F/R removed the CarR ORF from pcarR-3F, and the product was ligated via Gibson assembly to create p*3F*.
ii. pMCSG53-*carR*. To express CarR, primers EX-carR-F and EX-carR-R were used to PCR amplify the whole coding region of CarR. The resulting amplicon was assembled into the pMCSG53 backbone and prepared by PCR using the primer set pMCSG53(5kb)-F and pMCSG53(5kb)-R. The recombinant plasmid was then introduced into *E. coli* BL21(DE3). The His_6_-tagged CarR protein was purified by affinity chromatography following a published protocol (40). The purified protein was subsequently used for EMSA experiments and for generating polyclonal antibodies through Cocalico Biologicals, Inc.
iii. pCWU6-*pfruR(P1)-luc* & pCWU6-*pfruR(P2)-luc*. The primer pair pfruR-F/R was used to amplify the promoter region of *fruR*. The luciola red luciferase gene (*luc*) was PCR-amplified from pBCG06 (41) using the primers luc-F/R. The *pfruR* amplicon was then cloned upstream of *luc* via Gibson assembly, using KpnI/HindIII-digested pCWU6 to create pCWU6-*pfruR(P1)-luc*. To introduce mutations in the putative CarR binding sites within the *fruR* promoter, we incorporated the mutation site in the primer pfruR-R2. We used it with pfruR-F to amplify the mutated *fruR* promoter. The mutated promoter was then ligated with *luc* and integrated into digested pCWU6.
iv. p*Hp*. A fragment containing the promoter region of the putative hypothetical gene (*hp*, C4N14_10620) and its operon reading frame was PCR-amplified using primers pfruR-F and hp-R. The *hp* amplicon was cloned into KpnI/HindIII-digested pCWU6 to create p*Hp*.
v. p*radD*. We performed an inverse PCR with primers (galKrem-F/R) to remove *galKca* from pZP05, generating the shuttle plasmid pBCG11. The primer pair Pfdx-F/R was used to PCR amplify the promoter region of *fdx* from plasmid pZP06 (37), while primers com-radD-F/R targeted the *radD* open reading frame. Both amplicons were then cloned into the pBCG11 backbone, which was linearized by PCR using primers pBCG11-F/R.
vi. pP*radA*-*luc* The primer pair pradA-F/R was used to amplify the promoter region of *radABCD*. The luciola red luciferase gene (*luc*) was PCR-amplified from pBCG06 (41) using the primers luc-F2/R. The *pradA* amplicon was then cloned upstream of *luc* via Gibson assembly, using pCWU6 as the vector backbone, which had been linearized with *KpnI* and *HindIII* digestion. The resulting plasmid, pP*radA-luc*, was introduced into the wild-type *Fusobacterium nucleatum* strain ATCC 23826, creating a reporter strain for monitoring *radABCD* expression.
vii. p*fruB* The primer pair PfruR-F/PfruR3 was used to amplify the promoter region of the *fruRBA* operon, while the *fruB* gene was PCR-amplified from chromosomal DNA using primers com-fruB-F/com-fruB-R. The amplified *pfruR* fragment was cloned upstream of *fruB* via Gibson assembly, using pCWU6 as the vector backbone.

### Western blotting analysis

Various *Fusobacterium* strains were grown to late log phase with TSPC media in an anaerobic chamber, and cells from 1 mL of each culture were collected. The cell pellets were washed twice with water and resuspended in sodium dodecyl sulfate (SDS) sample buffer. Samples were boiled for 10 minutes (for detection of RadD, samples were heated at 70°C for 10 min), and proteins were separated on a 4%–20% Tris-glycine gradient SDS-PAGE (BIO-RAD, #456109), transferred to a PVDF membrane, and probed with rabbit anti-RadD (1:1000 dilution), rabbit anti-FLAG M2 antibody (#2368, Cell Signaling Technology; 1:3000 dilution), and anti-FomA (1:1000 dilution) as previously used (42). FomA, an outer membrane protein (OMP), was a loading control. Polyclonal goat anti-rabbit IgG conjugated with IRDye 680LT (#92668021; LI-COR Biosciences, USA) was used to detect the primary antibodies at a 1:5000 dilution.

### Bacterial co-aggregation

Co-aggregation assays were performed with *F. nucleatum* wild-type and its derivatives and *A. oris* MG-1, as previously described (43). Briefly, stationary-phase cultures of bacterial strains were grown in TSPC or heart infusion broth (for MG-1) harvested by centrifugation, washed in a NaN_3_-free coaggregation buffer (0.1 mM CaCl_2_, 0.1 mM MgCl_2_, 0.15 M NaCl and 1 mM Tris buffer adjusted to pH8.0) (44), and suspended to an equal cell density of approximately 2 x 10^9^ ml^-1^ based upon OD_600_ values. For coaggregation, 0.2 mL aliquots of *Actinomyces* and *Fusobacterium* cell suspensions were combined in a 24-well plate, mixed on a rotating shaker for several minutes, and imaged.

### Quantitative real-time PCR

To assess whether the 3xFLAG-tag affects CarR function, we examined the wild-type strain Δ*carR*, and its derivatives containing p*carR*, p*carR-3F*, or p*3F*. Each strain was initiated from a single colony and inoculated into a TSPC medium, with and without 5 µg/ml thiamphenicol. Following overnight growth, each culture was subcultured at a 1:20 dilution into fresh medium and allowed to grow to an OD₆₀₀ of approximately 0.8 (mid-llog phase).

To determine whether coaggregation serves as a signal sensed by the two-component system CarRS, we examined *rad* gene operon expression in *F. nucleatum* strains WT, Δ*radD*, Δ*carS*, and Δ*carS* p*radD*. Each *Fusobacterium* strain and *A. oris* MG-1 or S. oralis #34 was cultured for 16 hours to reach the stationary phase, with OD₆₀₀ adjusted to 1.0 before single-strain and co-culture assays. For single-strain assays, 8 mL of each OD₆₀₀-adjusted culture was centrifuged, the pellet was washed with sterile H₂O, resuspended in 8 mL of fresh TSCP medium, and incubated anaerobically for 2 hours before RNA extraction. In co-culture assays, three conditions were tested: (1) Simple Co-Culture Mix: 8 mL each of *Fusobacterium* and *A. oris* MG-1 cultures were centrifuged, washed with sterile H₂O, resuspended in 4 mL of TSCP medium, combined to a final volume of 8 mL, and incubated anaerobically for 2 hours; (2) High-Density Co-Culture: the simple co-culture mix was further centrifuged at 6,000 rpm for 5 minutes to form a high-density pellet, which was left intact and incubated anaerobically for 2 hours; and (3) Coaggregation Clump Formation: *F. nucleatum* and *A. oris* MG-1 cell pellets (from 8 mL cultures) were washed with coaggregation buffer, resuspended in 1 mL, and gently mixed to encourage visible coaggregation clumps. After allowing clumps to settle at the tube bottom (∼10 minutes), the supernatant was carefully removed, and 8 mL of TSPC medium was added without disrupting the clumps, followed by anaerobic incubation for 2 hours.

For both assays, cells from each condition were harvested by centrifugation, resuspended in 1 mL Trizol (Ambion), and lysed through mechanical disruption with 0.1-mm silicon beads (MP Bio). Total RNA was extracted using the Direct-zol RNA MiniPrep kit (Zymo Research), and cDNA synthesis was performed with SuperScript III reverse transcriptase (Invitrogen). Quantitative PCR was then conducted using iTaq SYBR Green Supermix (Bio-Rad) with primers targeting *radD*, *fruA*, *radB*, and *fruR* (Table S2). Gene expression levels were calculated using the 2^–ΔΔCt method, normalized to the *gyrB* gene. All experiments were validated in two independent runs, each performed in triplicate.

### ChIP-seq and bioinformatics analysis

The p*carR-3F* plasmid encoding CarR-FLAG was introduced into the Δ*carR* strain for ChIP assays with modifications to an established protocol (25). The Δ*carR*p*carR-3F* strain was grown overnight in TSPC medium, diluted 1:20 into fresh medium, and cultured to an OD_600_ of 0.8. Crosslinking was performed with 1% formaldehyde for 20 minutes at room temperature, followed by quenching with 0.5 M glycine. Cells were collected, washed with ice-cold PBS, and lysed by French Press in IP buffer [50 mM HEPES, 150 mM NaCl, 1 mM EDTA] with protease inhibitors. DNA was sheared to 100-500 bp using Bioruptor Pico®, and the supernatant was used as input for immunoprecipitation (IP) after removing cellular debris by centrifugation. The IP sample was incubated overnight with 35 µl Anti-FLAG M2 Magnetic Beads (Sigma, M8823), washed with low-salt (10 mM Tris-HCl [pH8.0], 1 mM EDTA, and 150 mM NaCl), high-salt (10 mM Tris-HCl [pH8.0], 1 mM EDTA and 500 mM NaCl), and TE buffers (10 mM Tris-HCl [pH8.0], 1 mM EDTA), and eluted in buffer (50 mM Tris-HCl [pH8.0], 10 mM EDTA, 1% SDS) with 5 M NaCl at 65°C for 12 hours to reverse crosslinks. Following RNaseA treatment, DNA was purified by phenol-chloroform extraction. For library preparation, 100-300 bp fragments were selected with Kapa Pure Beads. Library construction involved end repair, A-tailing, adapter ligation, and size selection, followed by quantification and quality assessment using an Agilent Bioanalyzer 2100 and qPCR. Libraries were sequenced using a paired-end 75-cycle on an Illumina NextSeq 550 at the Cancer Genomics Center at The University of Texas Health Science Center at Houston (CPRIT RP240610). Paired-end reads from two independent ChIP-seq experiments were aligned to the *F. nucleatum* subsp. *nucleatum* ATCC 23726 genome (NCBI RefSeq assembly: GCF_003019785.1) using BWA. Peaks enriched in ChIP-seq reads (q ≤ 0.05) were identified with MACS2 (45), visually inspected with IGV, and normalized by coverage per base. The CarR-binding consensus motif was generated using MEME (46).

### Identification of transcription start site by 5′ RACE

To identify the transcription start site of the *radABCD* transcript, 5′ random amplification of cDNA ends (5′ RACE) was performed using the FirstChoice RLM RACE kit (Invitrogen™, #AM1700M) following the manufacturer’s instructions. For the first PCR round, the 5′ RACE outer primer and the *radB* outer reverse primer (Table S2) were used with reverse transcription products as the template. The first-round PCR products were then subjected to a second round of amplification using the 5′ RACE inner primer and the *radB* inner reverse primer (Table S2). The resulting 5′ RACE products were cloned into the pCM-*galk* vector (47) via Gibson assembly and sequenced with the S-pCM-galK-F primer (Table S2).

### Electrophoretic Mobility Shift Assays (EMSA)

For EMSA studies, Cy5-labeled promoter regions of *radA*, *fruR*, *megL*, *gcdD*, *kal*, and *fap2* were amplified using specific primers (Table S2). These probe fragments were incubated for 20 minutes at room temperature with various concentrations of purified CarR-His₆ in a 20 µl binding reaction buffer consisting of water, 1x binding buffer, 50 ng/µl poly (dI-dC), 2.5% glycerol, 5 mM MgCl₂, and unlabeled specific probe as needed, following the LightShift Chemiluminescent EMSA kit protocol (Pierce, #20158). The samples were then loaded onto a 6% polyacrylamide gel and electrophoresed at 100 V for 1 hour at room temperature. Gels were imaged using the ChemiDoc System (Bio-Rad). For competitive EMSA, probes P1, P2, P3, and P4 (Table S2) were synthesized by Twist Bioscience.

### DNase I Footprinting assay

The DNase I footprinting assay, based on dye/primer sequencing, was performed following a previously described protocol (25). Briefly, the *radA* promoter region was PCR-amplified with *pfu* DNA polymerase using primers pradA Footprinting FAM F (5’-end labeled with 6-carboxyfluorescein [6-FAM]) and pradA FAM R (Table S2). The FAM-labeled probes were purified and quantified with an Implen NanoPhotometer® N50. For each assay, 200 ng of the labeled probe was incubated with 0 or 7 µM CarR in a 20 µl binding buffer for 30 minutes at 25°C. Then, 0.015 U DNase I (Thermo Scientific) was added, and the mixture was further incubated for 1 minute at 25°C for fragmenting DNA. The reaction was stopped by adding 2.77 µl of 5 mM EDTA and heating at 65°C for 10 minutes. The digested DNA fragments were purified using the Agencourt AMPure XP PCR Purification Kit (Beckman Coulter) and sent to GENEWIZ for fragment analysis. For analysis, 1 µl of digested DNA was mixed with 8.5 µl of highly deionized formamide and 0.5 µl of GeneScan™ LIZ500/LIZ200 Size Standard (Applied Biosystems). Samples were analyzed on an ABI 3730xl DNA Analyzer (Applied Biosystems), and results were processed with Peak Scanner Software (Applied Biosystems).

### Gene deletion in *F. nucleatum*

To create deletion constructs for the hypothetical gene *hp* (C4N14_10620), located just upstream of *fruB*, 1.0-kb fragments flanking the region encoding amino acids 2-31 were PCR-amplified with specific primers (Table S2). For the *fruB* deletion plasmid, 1.0-kb regions upstream and downstream of *fruR* were amplified using fruBupF/R and fruBdnF/R primers. Overlapping PCR was used to fuse each upstream and downstream fragment, and the fused segments were cloned into pBCG02 (48). The resulting plasmids were introduced into *F. nucleatum* ATCC 23726 by electroporation, and in-frame deletion mutants were selected using the *hicA* counterselection marker (48).

The generation of a nonpolar, in-frame *fruB* deletion mutant in *F. nucleatum* was performed following a previously published protocol (32). Briefly, 1-kb fragments upstream and downstream of the *fruB* gene were cloned into the integrative vector pCM-GalK, which contains genes conferring thiamphenicol resistance and *galK* expression (32) The resulting plasmid was electroporated into *F. nucleatum* CW1, a strain lacking a functional *galK* gene. Integration of the vector into the bacterial chromosome via homologous recombination was selected on TSPC agar plates containing 5 μg/mL thiamphenicol. Resolution of the vector through a second homologous recombination event, leading to either deletion of the target gene or reconstitution of the wild-type genotype, was selected on plates containing 0.2% 2-deoxygalactose (2-DG). Colonies were resistant to 2-DG and sensitive to thiamphenicol were screened by PCR to confirm the expected *fruB* deletion.

### Growth assay

*F. nucleatum* wild-type and derived strains were cultured overnight, starting with 2-3 single colonies on the plates. These overnight cultures were then diluted 1:20 into fresh TSPC medium, with and without 20 mM fructose. After 16 hours, when the cultures reached the stationary phase, the OD_600_ was measured and recorded using an Implen OD600® device. The experiment was repeated at least three times.

### Luciferase Assays

Overnight cultures of *F. nucleatum* wild-type, Δ*carR*, and Δ*carS* strains carrying either the pCWU6-*PfruR(P1)-luc* or pCWU6-*PfruR(P2)-luc* vector were sub-cultured 1:20 in TSPC medium. Once the cultures reached an OD_600_ of approximately 1, three 100 µL aliquots from each culture were transferred into a 96-well microplate (Corning no. 3917). Each aliquot was mixed with 25 µL of 1 mM d-luciferin (Molecular Probe) in 100 mM citrate buffer (pH 6.0) and vigorously pipetted 10 times under aerobic conditions. Luciferase activity was measured with a GloMax Navigator Microplate Luminometer, and culture growth was assessed by measuring OD_600_. As a negative control, the wild-type strain carrying an empty pCWU6 plasmid showed minimal background signal, which was subtracted from the test samples. Each experiment was performed in triplicate for accuracy.

## AUTHOR CONTRIBUTIONS

C. W. conceived and supervised the project. B.G. and C. W. designed the project. B.G. and C.W. performed all the experiments. B.G. and C. W. analyzed data. C.W. and B. G. wrote the manuscript with contribution and approval from all authors.

## DATA AVAILABILITY

The ChIP-seq data has been deposited in the NCBI SRA under accession number SUB14833047. All additional study data are provided within the article and/or supplementary materials.

## ACKNOWLEDGMENTS

This research received funding from the National Institute of Dental and Craniofacial Research (NIDCR) under grant number DE030895, awarded to C.W. We thank the technical support from the Cancer Prevention and Research Institute of Texas (CPRIT RP240610)

